# Theoretical Analysis of Inducer and Operator Binding for Cyclic-AMP Receptor Protein Mutants

**DOI:** 10.1101/236455

**Authors:** Tal Einav, Julia Duque, Rob Phillips

## Abstract

Allosteric transcription factors undergo binding events both at their inducer binding sites as well as at distinct DNA binding domains, and it is often difficult to disentangle the structural and functional consequences of these two classes of interactions. In this work, we compare the ability of two statistical mechanical models – the Monod-Wyman-Changeux (MWC) and the Koshland-Némethy-Filmer (KNF) models of protein conformational change – to characterize the multi-step activation mechanism of the broadly acting cyclic-AMP receptor protein (CRP). We first consider the allosteric transition resulting from cyclic-AMP binding to CRP, then analyze how CRP binds to its operator, and finally investigate the ability of CRP to activate gene expression. In light of these models, we examine data from a beautiful recent experiment that created a single-chain version of the CRP homodimer, thereby enabling each subunit to be mutated separately. Using this construct, six mutants were created using all possible combinations of the wild type subunit, a D53H mutant subunit, and an S62F mutant subunit. We demonstrate that both the MWC and KNF models can explain the behavior of all six mutants using a small, self-consistent set of parameters. In comparing the results, we find that the MWC model slightly outperforms the KNF model in the quality of its fits, but more importantly the parameters inferred by the MWC model are more in line with structural knowledge of CRP. In addition, we discuss how the conceptual framework developed here for CRP enables us to not merely analyze data retrospectively, but has the predictive power to determine how combinations of mutations will interact, how double mutants will behave, and how each construct would regulate gene expression.

## Introduction

Cyclic-AMP receptor protein (CRP; also known as the catabolite receptor protein, CAP) is an allosteric transcription factor that regulates over 100 genes in *Escherichia coli* (1–4). Upon binding to cyclic-AMP (cAMP), the homodimeric CRP undergoes a conformational change whereby two alpha helices reorient to open a DNA binding domain (5), allowing CRP to bind to DNA and affect transcription (6–8). While much is known about the molecular details of CRP and how different mutations modify its functionality (9, 10), each new CRP mutant is routinely analyzed in isolation using phenomenological models. We argue that given the hard-won structural insights into the conformational changes of proteins like CRP, it is important to test how well mechanistically motivated models of such proteins can characterize the wealth of available data.

One of the difficulties inherent in understanding allosteric transcription factors such as CRP lies in our inability to disentangle the numerous processes involved in transcription, such as the binding of cAMP to CRP, of CRP to DNA, and of transcription regulation as shown in Fig. 1(A). Measurements of gene expression depend upon all of these processes, and mechanistic statements about the individual steps must necessarily be inferred (11–14). To this end, *in vitro* studies are beginning to probe each binding event separately, providing a testbed to refine our understanding of both allostery and transcriptional activation.

**Figure 1.**
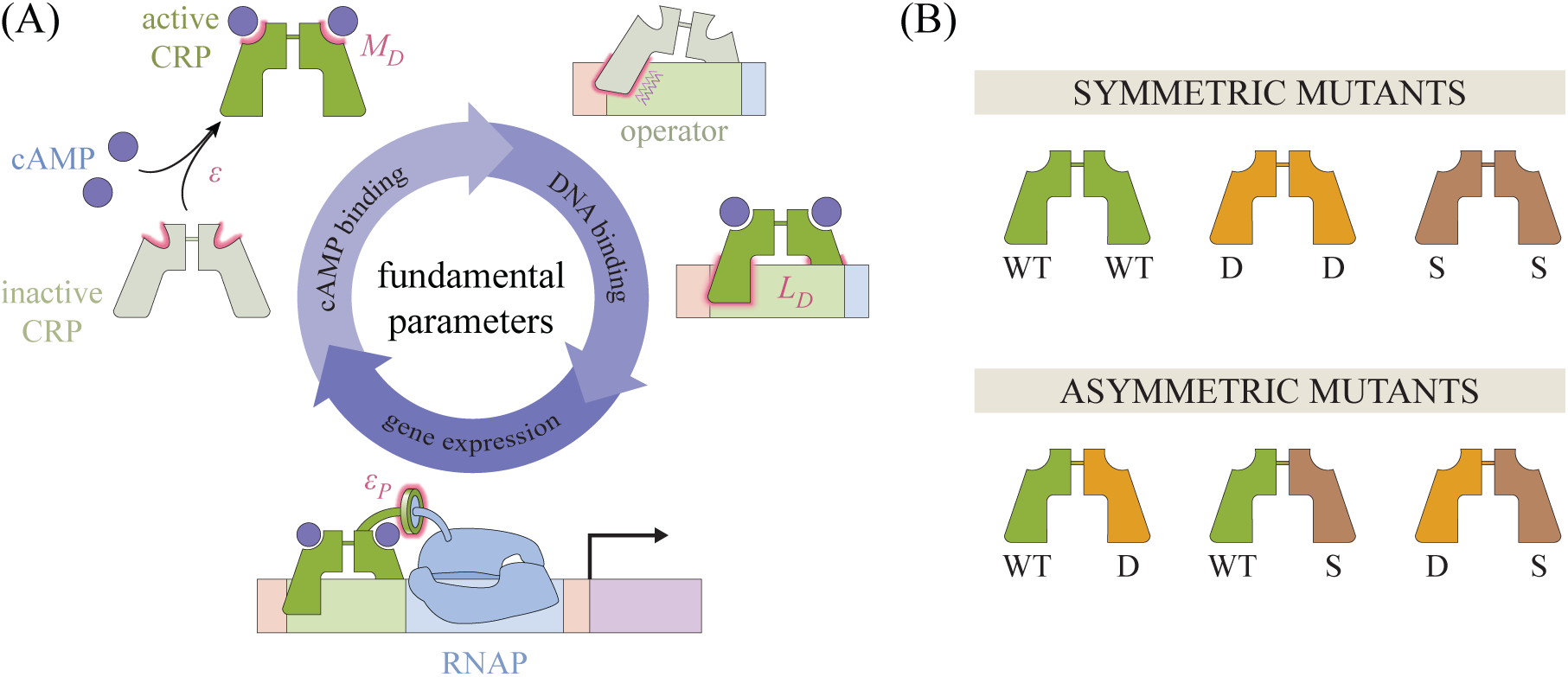
Key parameters governing CRP function. (A) Within the MWC and KNF models, each CRP subunit can assume either an active or an inactive conformation with a free energy difference *є* between the two states. cAMP can bind to CRP (with a dissociation constant 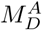 in the active state and 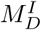 in the inactive state) and promotes the active state (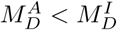 in the MWC model; 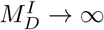 in the KNF model). Active CRP has a higher affinity for the operator 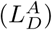 than the inactive state 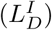. When CRP is bound to DNA, it promotes RNA polymerase binding through an interaction energy *є*_*P*_, thereby enhancing gene expression. (B) Lanfranco *et al.* constructed a single-chain CRP molecule whose two subunits could be mutated independently. They measured the cAMP and DNA binding affinity for CRP mutants comprised of wild type (WT), D (D53H), or S (S62F) subunits.

Our paper is inspired by a recent *in vitro* study of CRP performed by Lanfranco *et al.* who separately measured the two binding events of CRP, first to cAMP and then to DNA (15), providing an opportunity to make a rigorous, quantitative analysis of the allosteric properties of CRP. To this end, we explore two mechanistic frameworks for the allosteric transition of CRP: the Monod-Wyman-Changeux (MWC) model, which posits that both CRP subunits fluctuate concurrently between an active and inactive conformational state (16), and the Koshland-Némethy-Filmer (KNF) model, which proposes that each subunit must independently transition from an inactive to active state upon ligand binding (17). Although the MWC model provides a marginally better characterization of the data, the two models offer different interpretations of the general behavior of CRP. For example, the parameter set we inferred for the KNF model predicts that CRP will bind equally well to DNA regardless of whether it is bound to one or two cAMP molecules, while the MWC model predicts that singly bound CRP will bind more tightly to DNA than unbound and doubly bound CRP. Knowledge of the structure of CRP is in line with the MWC predictions (10, 18), demonstrating that a model should not be judged merely by its goodness of fit, but rather by the interpretation of its fit parameters compared with the available knowledge of the system.

In addition to their consideration of the wild type protein, Lanfranco *et al.* engineered a single-chain CRP molecule whose two subunits are tethered together by an unstructured polypeptide linker. This construct enabled them to mutate each subunit independently as shown in Fig. 1(B), providing a novel setting within which to analyze the combinatorial effects of mutations. Specifically, they took three distinct CRP subunits – the wild type subunit and the well characterized mutations D53H and S62F originally chosen to perturb the transcription factor’s cAMP binding domain (19, 20) – and linked them together in every possible combination to create six CRP mutants.

The effects of mutations are often difficult to interpret, and indeed the results from Lanfranco *et al.* showed no clear pattern. The behavior of each mutant was analyzed independently by fitting its binding curve to a second order polynomial, but this analysis was unable to make use of the fact that the six CRP mutants are linked through their subunit compositions (15). In this work, we aim to close that gap by constructing a quantitative framework that can describe the full suite of CRP data by utilizing the subunit composition of CRP.

This concrete link between the composition and behavior of CRP mutants raises the question of whether the response of a mutant can be predicted based on the behavior of closely related mutants. For example, given sufficient data of CRP with two wild type (WT) subunits and of CRP with two D53H subunits, can we predict how a CRP comprised of one wild type and one D53H subunit will behave? More generally, can the behavior of the symmetric mutants (top row of Fig. 1(B)) predict the behavior of the asymmetric mutants (bottom row of Fig. 1(B))? We demonstrate that both the MWC and KNF models can generate such predictions. These results suggest a way to harness the combinatorial complexity of oligomeric proteins and present a possible step towards systematically probing the space of mutations.

We end by exploring the physiological impact of these CRP mutants by considering how they would promote gene expression *in vivo.* Because CRP is a global activator, its activity within the cell is tightly regulated by enzymes that produce, degrade, and actively transport cAMP. We discuss how these processes can either be modeled theoretically or excised experimentally and calibrate our resulting framework for transcription using gene expression measurements for wild type CRP (7). In this manner, we find a small, self-consistent set of parameters able to characterize each step of CRP activation shown in Fig. 1(A).

The remainder of this paper is organized as follows. First, we characterize the interaction between cAMP and CRP for the six CRP mutants created by Lanfranco *et al.* and quantify the key parameters governing this behavior. Next, we analyze the interaction between CRP and DNA from the perspectives of the MWC and KNF models and discuss how the interpretations of the parameters inferred by the MWC model are more in line with structural knowledge of the system. Finally, we consider how CRP enhances gene expression and extend the results from Lanfranco *et al.* to predict the activation profiles of the CRP mutants within a cellular environment.

## Results and Discussion

### The Interaction between CRP and cAMP

In this section, we examine the cAMP-CRP binding process through the lenses of the MWC and KNF models. We find that both models can characterize data from a suite of CRP mutants using a compact set of parameters, thereby highlighting how each mutant’s behavior is tied to its subunit composition.

#### MWC Model

We first formulate a description of cAMP-CRP binding using the MWC model, where the two subunits of each CRP molecule fluctuate concurrently between an active and inactive state. We define the free energy difference between the inactive and active conformations to be є per subunit, so that the total free energy difference between inactive CRP and active CRP is 2*є*. The different conformations of CRP binding to cAMP and their corresponding Boltzmann weights are shown in Fig. 2(A). For each cAMP-CRP dissociation constant 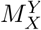, the subscript denotes which CRP subunit it describes – either the left (*L*) or right (*R*) subunit – while the superscript denotes the active (*A*) or inactive (*I*) state of CRP. Given a cAMP concentration [*M*], the fraction of occupied cAMP binding sites is given by

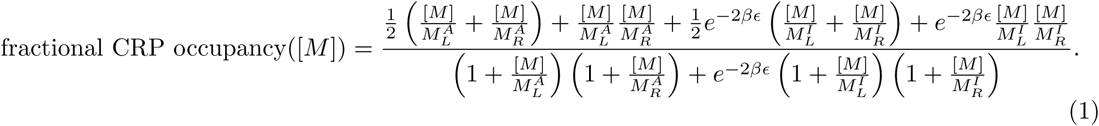

**Figure 2.**
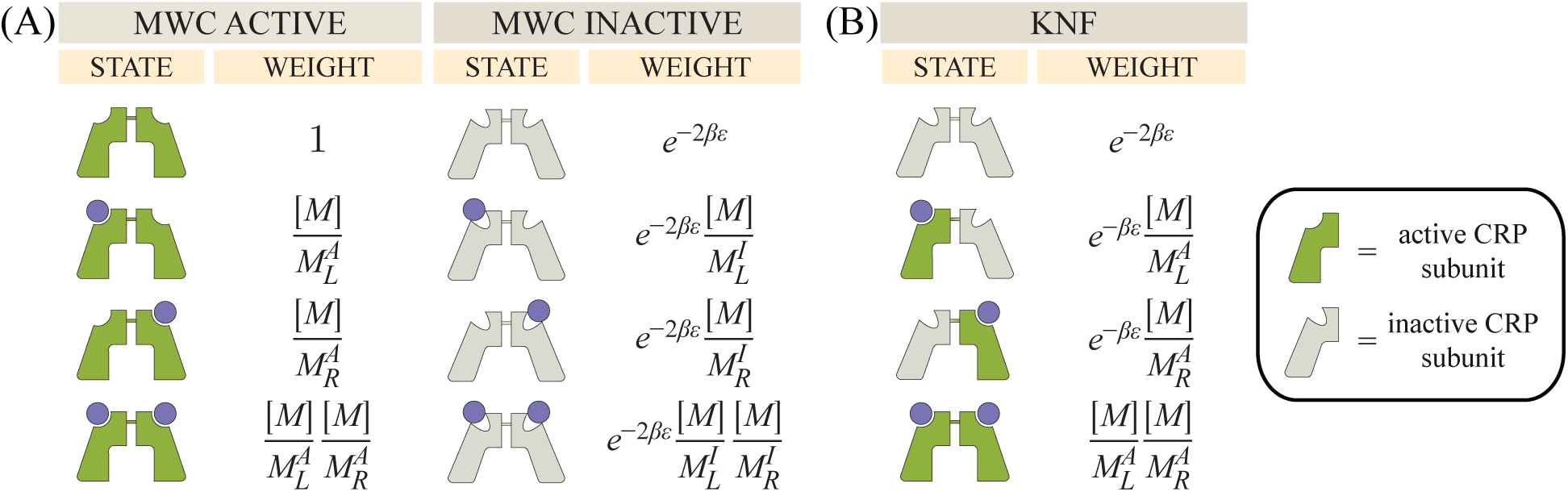
Macroscopic states and Boltzmann weights for cAMP binding to CRP. (A) Within the MWC model, cAMP (purple circles) may bind to a CRP subunit in either the active (dark green) or inactive (light green) state. 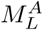 and 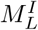 represent the dissociation constants of the left subunit in the active and inactive states, respectively, while 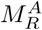 and 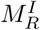 represent the analogous dissociation constants for the right subunit. [*M*] denotes the concentration of cAMP and є represents the free energy difference between each subunit’s inactive and active states. (B) The KNF model assumes that the two CRP subunits are inactive when unbound to cAMP and transition to the active state immediately upon binding to cAMP. The parameters have the same meaning as in the MWC model, but states where one subunit is active while the other is inactive are allowed.

Here, the fractional occupancy of CRP bound to zero, one, or two cAMP equals 0, ½, and 1, respectively. Experimentally, the fractional occupancy was measured using ANS fluorescence which utilizes a fluorescent probe triggered by the conformational change of cAMP binding to CRP (15). We note that this measurement was carried out *in vitro* in the absence of DNA.

Lanfranco *et al.* considered CRP subunits with either the D53H or S62F point mutations (hereafter denoted by D and S, respectively), with the D subunit binding more strongly to cAMP than the wild type while the S subunit binds more weakly as shown in Fig. 3(A). While we could characterize the dose-response curves of each CRP mutant independently – for example, by using Eq. 1 to extract a set of parameters for each mutant – such an analysis lacks a direct connection between the subunit composition and the corresponding binding behavior. Instead, we assume that the cAMP binding affinity for each subunit should be uniquely dictated by that subunit’s identity as either the WT, D, or S subunit. To that end, we represent the fractional occupancy of CRP_D_/_WT_ using Eq. 1 with one D subunit 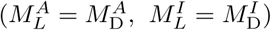 and one WT subunit 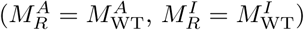. The equations for the remaining CRP mutants follow analogously, tying the behavior of each mutant to its subunit composition.

**Figure 3.**
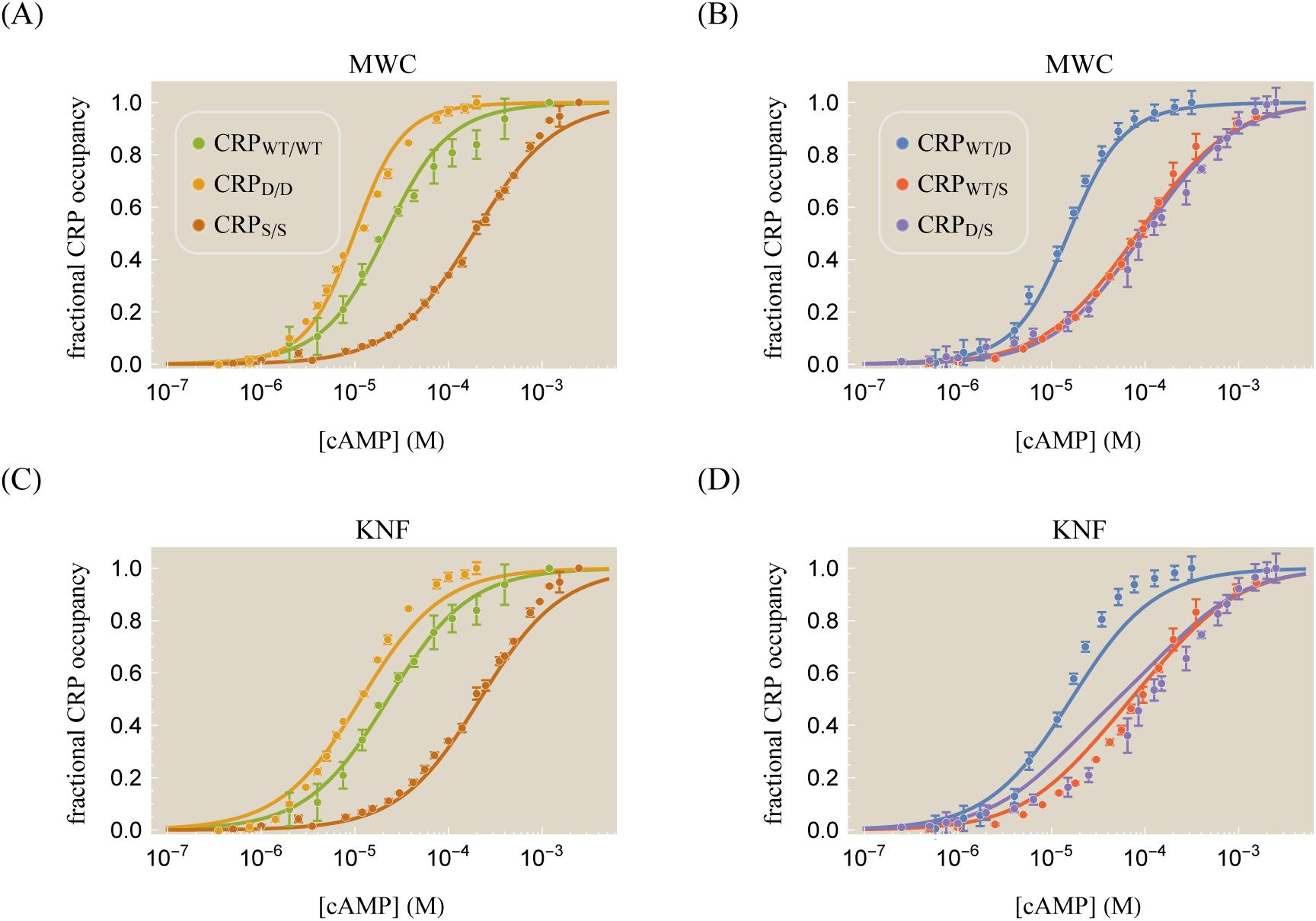
cAMP binding curves for different CRP mutants. In addition to the wild type CRP subunit (denoted WT), the mutation D53H (denoted D) and the mutation S62F (denoted S) can be applied to either subunit as indicated by the subscripts in the legend. (A) Curves were characterized using the MWC model, Eq. 1. The D subunit increases CRP’s affinity for cAMP while the S subunit decreases this affinity. (B) Asymmetrically mutating the two subunits results in distinct cAMP binding curves. The data for the WT/D mutant lies between the WT/WT and D/D data in Panel (A), and analogous statements apply for the WT/S and D/S mutants. (C) The symmetric mutants and (D) the asymmetric mutants can also be analyzed using the KNF model, Eq. 6, resulting in curves that are similar to those found by the MWC model. The (corrected) sample standard deviation 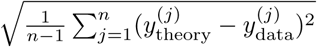 equals 0.03 for the MWC model and 0.06 for the KNF model, and the best-fit parameters for both models are given in Table 1.

**Table 1.**
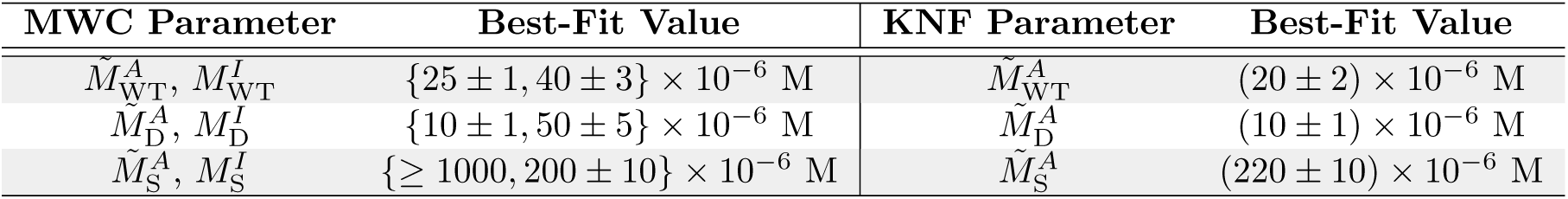
**Parameters for cAMP binding to CRP.** The data in Fig. 3 can be characterized using a single set of dissociation constants for the WT, D, and S subunits whose values and standard errors are shown. The left column corresponds to the MWC parameters given in Eq. 1 while the right column corresponds to the KNF model given by Eq. 6. For both models, the data only constrains the parameter combination 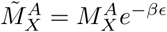. The effective dissociation constant of the S subunit in the MWC model can only be bounded from below as 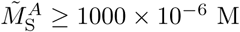.

Many studies have been conducted upon cAMP-CRP binding using a broad range of methods including equilibrium dialysis, fluorescence assays, and protease digestion, but the resulting parameters inferred from these experiments have varied widely (19). Estimates on the free energy difference 2*є* between inactive and active CRP range from approximately –20 *k_B_T* to –10 *k_B_T* (10, 21, 22) while apparent dissociation between cAMP and CRP_WT_/_WT_ range from 10^−6^–10^−3^ M (19, 23, 24).

Part of the difficulty in pinning down these values stems from the fact that degenerate parameter values reproduce equivalent binding curves. For instance, it is known that in the absence of cAMP, the overwhelming majority of CRP molecules will be in the inactive state (1 ≪ *e*^−2*β*_є_^). Intuitively, there will effectively be no active CRP molecules in the absence of cAMP regardless of whether 2*є* = —10 *k_B_T* or 2*є* = —20 *k_B_T*; however, in the presence of saturating cAMP the latter case will require a larger affinity between active CRP and cAMP (smaller 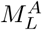 and 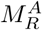) to compensate for this greater free energy difference. The same cAMP-CRP binding curves can even be produced for an arbitrarily large and negative free energy difference (*є* → —∞) provided that the dissociation constants scale appropriately. This scaling can be determined by fixing the ratio of doubly-cAMP bound CRP in the active state 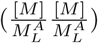 and inactive state 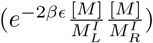. For the CRP system this requires that 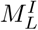 and 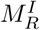 do not depend on *є* while 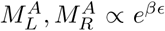. Mathematically, in the limit of a large and negative *є* the cAMP occupancy becomes

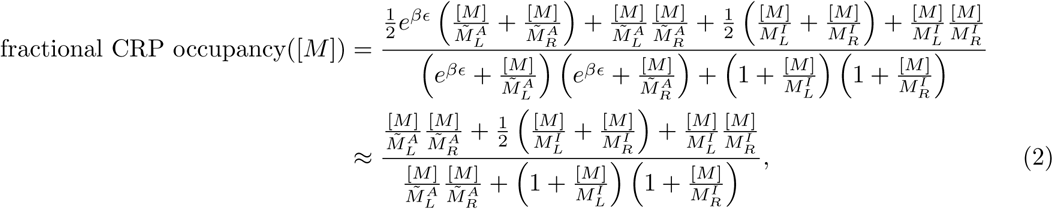

where in the first equality we multiplied the numerator and denominator of Eq. 1 by *e*^2*β*_є_^ and defined the effective dissociation constants

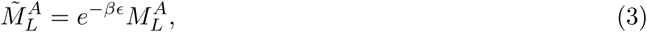

and

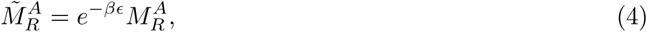

while in the latter equality of Eq. 2 we neglected all of the terms multiplied by the small quantity *e*^*β*_є_^ (which is equivalent to taking the zeroth order Taylor series about *e*^−*β*_є_^ ≈ 0). In Supporting Information Section A, we demonstrate how the e parameter may be shifted arbitrarily without altering the cAMP-CRP binding curves provided Eqs. 3 and 4 hold and that 2*є* ≲ —3 *k_B_T* (above which the approximation 1 ≪ *e*^−2*β*_*є*_^ breaks down). This last assumption is well justified, since the overwhelming majority of CRP molecules assume the inactive conformation in the absence of cAMP (10). Therefore, the maximum information that can be extracted from the data includes the inactive CRP-cAMP dissociation constants 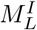 and 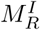 and the effective dissociation constants 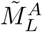 and 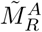. Lastly, we note that if the D and S mutations alter the free energy *є*, that effect will be absorbed into the effective dissociation constants.

Using Eq. 2, we can extract the set of effective dissociation constants for the WT, D, and S subunits that determine the behavior of all six CRP mutants. The resulting parameters (shown in Table 1) give rise to the cAMP-CRP binding curves in Fig. 3(A) and (B). In Supporting Information Section B, we demonstrate that the symmetric CRP mutants in Fig. 3(A) provide sufficient information to predict the behavior of the asymmetric mutants in Fig. 3(B). We further show that fitting each CRP data set individually to the MWC or KNF models without constraining the WT, D, and S subunits to a single unified set of dissociation constants results in only a marginal improvement over the constrained fitting. Finally, we analyze the slope of each cAMP binding response and explain why they are nearly identical for the six CRP mutants. Supporting Information Section C investigates the effects of the double mutation D+S on a single subunit.

#### KNF Model

We now turn to a KNF analysis of CRP, where the two subunits are individually inactive when not bound to cAMP and become active upon binding as shown in Fig. 2(B). Some studies have claimed that cAMP binding to one CRP subunit does not affect the state of the other subunit, in support of the KNF model (25). Other studies, meanwhile, have reported that a fraction of CRP molecules are active even in the absence of cAMP, thereby favoring an MWC interpretation (9). It is not yet known whether either model can accurately represent the system. To that end, we explore some of the consequences of a KNF interpretation of CRP.

Using the statistical mechanical states of the system in Fig. 2(B), the occupancy of CRP is given by

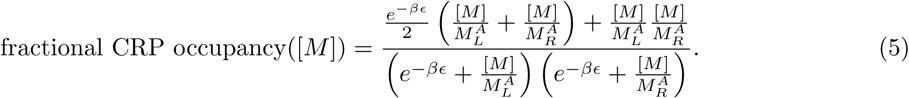

Note that by multiplying the numerator and denominator by *e*^2*β*_є_^ and defining the same effective dissociation constants (Eqs. 3 and 4) as for the MWC model, we can eliminate the free energy difference *є* to obtain the form

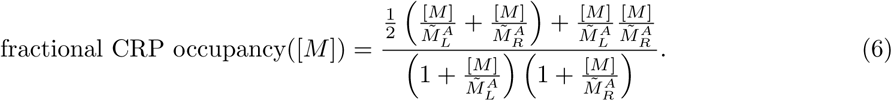

This simplification occurs because within the KNF model, a CRP monomer only switches from the inactive to active state upon cAMP binding. As a result, the free energy of cAMP binding to CRP and the free energy of the CRP undergoing its inactive-to-active state conformational always occur concurrently and may be combined into the effective dissociation constants 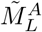 and 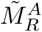. As shown in Fig. 3(C) and Fig. 3(D), the KNF model can characterize the six mutant CRP binding curves, albeit with a larger sample standard deviation than the MWC model. While the KNF model has fewer parameters, the cost of this simplicity is manifest in its slightly poorer fits.

### The Interaction between CRP and DNA

We now turn to the second binding interaction experienced by CRP, namely, that between CRP and DNA within the MWC and KNF models.

#### MWC Model

Consider a concentration [*L*] of CRP whose subunits either assume an active state (where they tightly bind to DNA with a dissociation constant *L_A_*) or in an inactive state (characterized by weaker DNA binding with dissociation constant *L_I_* satisfying *L_I_* > *L_A_*). The states and weights of this system within the MWC model are shown in Fig. 4(A).

**Figure 4.**
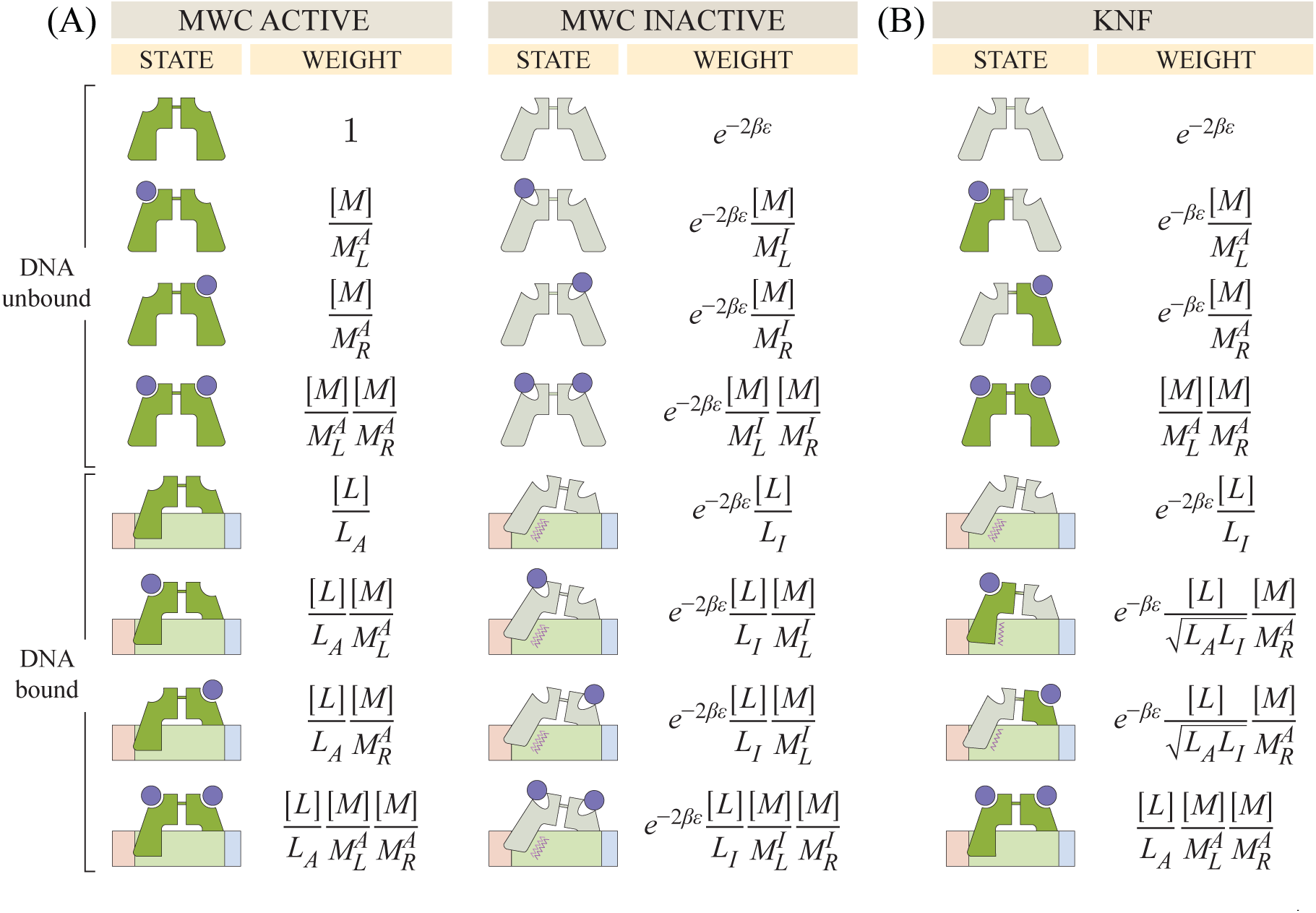
States and weights for CRP binding to DNA. (A) The DNA unbound states from Fig. 2 together with the DNA bound states. The Boltzmann weight of each DNA bound state is proportional to the concentration [*L*] of CRP and inversely proportional to the CRP-DNA dissociation constants *L_A_* or *L_I_* for the active and inactive states, respectively. (B) In the KNF model, these same parameters apply to each CRP subunit which can be independently active or inactive.

Lanfranco *et al.* fluorescently labeled a short, 32 bp DNA sequence which binds to CRP. Using a spectrometer, they measured the anisotropy of this fluorescence when different concentrations of CRP and cAMP were added *in vitro* (15). The data are shown in Fig. 5(A) for CRP_D/S_. When CRP binds, it slows the random tumbling of the DNA so that over very short time scales the fluorescence is oriented along a particular axis, resulting in a larger anisotropy readout. Unbound DNA is defined as having anisotropy = 1 while DNA-bound CRP with 0, 1, or 2 bound cAMP have higher anisotropies of 1 + *r*_0_, 1 + *r*_1_, and 1 + *r*_2_, respectively. Thus, the total anisotropy within the model is given by the weighted sum of each species (27), namely,

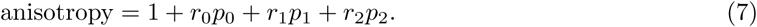

**Figure 5.**
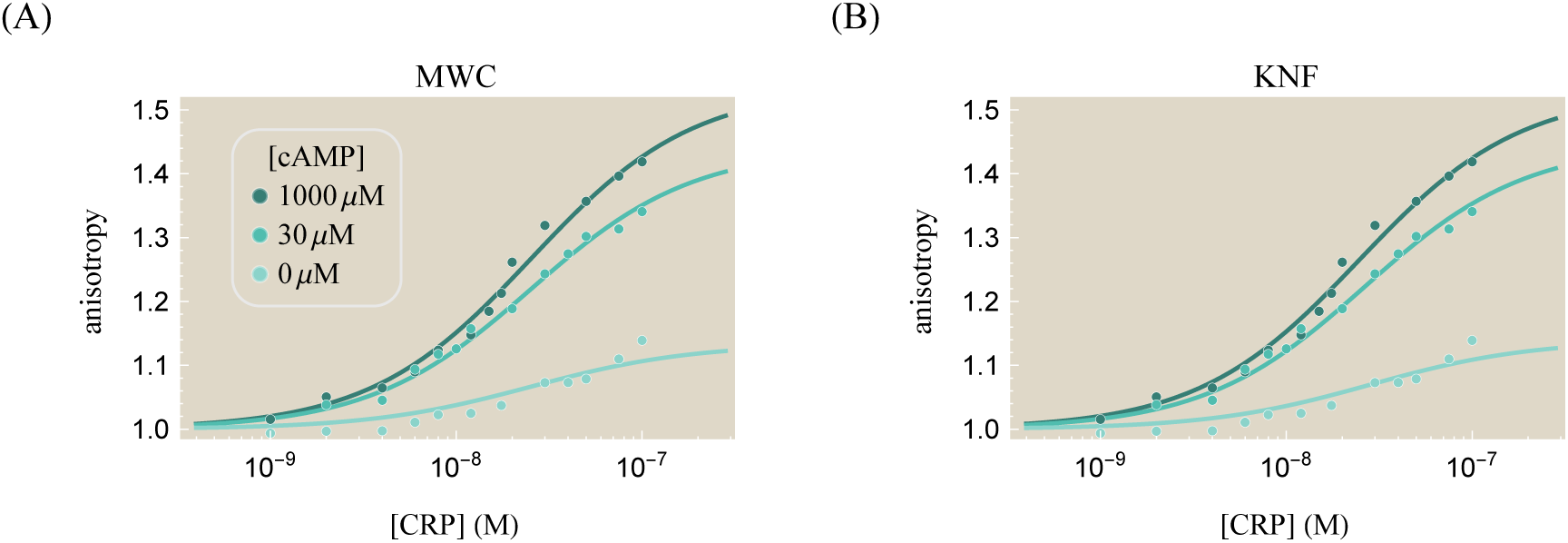
The interaction between CRP and DNA. (A) Anisotropy of 32-bp fluorescein-labeled *lac* promoter binding to CRP_D/S_ at different concentrations of cAMP. An anisotropy of 1 corresponds to unbound DNA while higher values imply that DNA is bound to CRP. (B) This same data analyzed using the KNF model. In the presence of cAMP, more CRP subunits will be active, and hence there will be greater anisotropy for any given concentration of CRP. The sample standard deviation 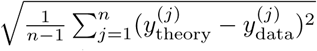 is 0.01 for both the MWC and KNF models, with the corresponding parameters given in Tables 1 and 2.

**Table 2.**
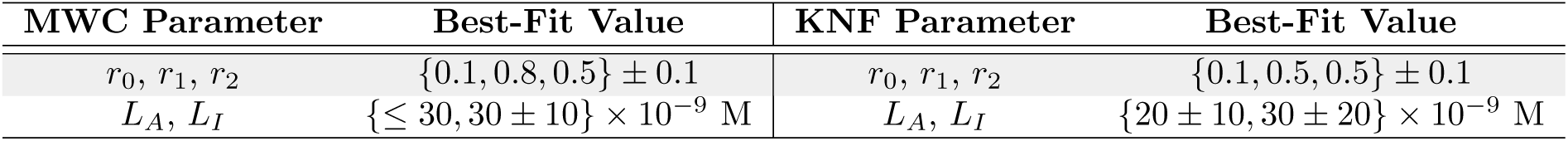
**Parameters for CRP binding to DNA.** The anisotropy data for CRP_D/S_ characterized using Eq. 7, as shown in Fig. 5. Each value is given as a mean ± standard error. The uncertainty in 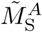 parameter (shown in Table 1) leads to a corresponding uncertainty in the active CRP dissociation constant *L_A_*.

| MWC Parameter | Best-Fit Value | KNF Parameter | Best-Fit Value |
| --- | --- | --- | --- |
| $r_0, r_1, r_2$ | $\{0.1, 0.8, 0.5\} \pm 0.1$ | $r_0, r_1, r_2$ | $\{0.1, 0.5, 0.5\} \pm 0.1$ |
| $L_A, L_I$ | $\{\leq 30, 30 \pm 10\} \times 10^{-9}$ M | $L_A, L_I$ | $\{20 \pm 10, 30 \pm 20\} \times 10^{-9}$ M |

Here, *p*_0_, *p*_1_, and *p*_2_ represent the probability that DNA-bound CRP will be bound to 0, 1, and 2 cAMP molecules, respectively. In this model, we have extended the classic MWC model to allow each of these states to have a unique DNA binding affinity. Using the effective dissociation constants (Eqs. 3 and 4) and neglecting all terms proportional to the small quantity *e*^*β*_*є*_^, we can write these probabilities as

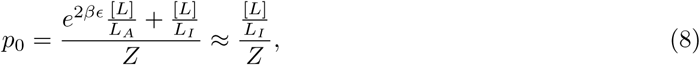

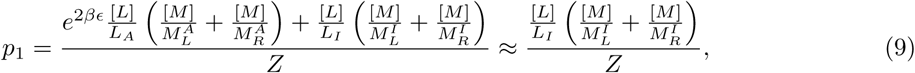

and

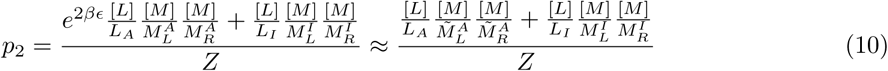

with

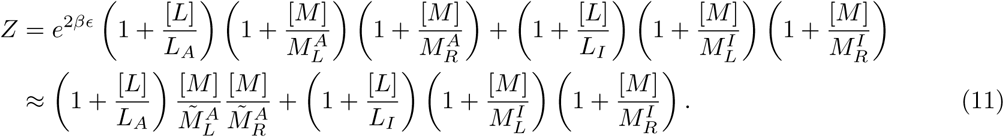

In making these approximations, we have assumed the stricter conditions 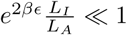 and 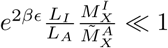 for the WT, D, and S subunits, all of which are valid assumptions for this system (see Supporting Information Section A).

Fig. 5(A) shows the resulting best-fit curves for the anisotropy data, with the corresponding CRP_D_/_S_ DNA dissociation constants given in Table 2. Since 1 + *r*_0_ ≈ 1, unbound CRP binds poorly to DNA, in accordance with the inactive state crystal structure whose DNA recognition helices are buried inside the protein (10). Additionally, the anisotropy 1 + *r*_1_ = 1.7 of the DNA-CRP-cAMP complex is larger than that of both the unbound state and the doubly bound state DNA-CRP-(cAMP)_2_ with 1 + *r*_2_ = 1.4; this suggests that CRP-(cAMP)_2_ binds more weakly to DNA that CRP-cAMP. Previous studies have confirmed this claim using multiple experimental methods including proteolytic digestion by subtilisin, chemical modification of Cys-178, and fluorescence measurements (18, 28), although other work has proposed an alternate explanation that above millimolar cAMP concentrations CRP attains new states with highly unfavorable DNA binding affinities (13, 29). In Supporting Information Section D, we extend the analysis of CRP-DNA binding to the remaining CRP mutants.

#### KNF Model

We now turn to the KNF model of DNA binding where each CRP subunit is inactive when not bound to cAMP and active when bound to cAMP as shown in Fig. 4(B). As in the MWC model, *L_A_* and *L_I_* represent the dissociation constants between DNA and CRP in the active and inactive states, respectively. The mixed state of DNA-bound CRP with one active and one inactive subunit has a dissociation constant 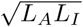 (see Supporting Information Section E). The anisotropy within the KNF model is given by Eq. 7 with

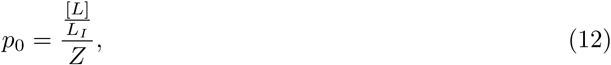

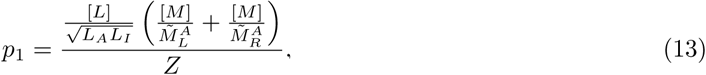

and

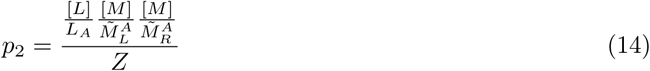

representing the probabilities of the DNA-CRP, DNA-CRP-cAMP, and DNA-CRP-(cAMP)_2_ states, respectively, with

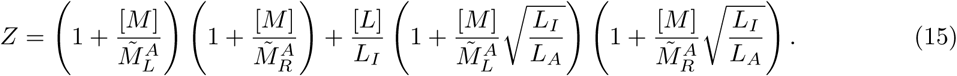

Fig. 5(B) shows the characterization of the anisotropy data using the KNF model with the corresponding parameters given by the right column of Table 2. Unlike the MWC model, the best-fit KNF anisotropy parameters satisfy *r*_1_ ≈ *r*_2_, suggesting that CRP binds equally tightly to DNA independent of how many cAMP are bound to it. Structural understanding of CRP support the MWC characterization, as both unbound and doubly bound CRP have been reported to have a significantly lower DNA-binding affinity than singly-cAMP bound CRP (18, 28).

### Implications of Mutations for *in vivo* Systems

CRP is a global transcriptional activator which governs many metabolic genes in *E. coli* (8). It is interesting to consider how the mutants characterized in the Lanfranco *et al.* experiments would behave as transcriptional activators for *in vivo* gene expression measurements. In this section, we make predictions for how CRP will act *in vivo* for the different mutants characterized above. To focus our analysis, we use the MWC model presented above to analyze gene expression measurements of a system where CRP is the only transcription factor present, although it is straightforward to generalize to more complex regulatory architectures or to apply the KNF framework to this process (30).

#### Simple Activation

As above, consider a cell with cAMP concentration [*M*] and CRP concentration [*L*] where the population of CRP is split between an active [*L_A_*] and an inactive [*L_I_*] conformation. Suppose the cell has a concentration [*P*] of RNA polymerase (RNAP) which have a dissociation constant *P_D_* with a promoter of interest. The thermodynamic states of the system are shown in Fig. 6, where the activator can bind to and recruit RNAP via an interaction energy *є*_*p*,*L_A_*_ between active CRP and RNAP with a weaker interaction *є*_*p*,*L_I_*_ between inactive CRP and RNAP. Without these two interaction energies, the RNAP and CRP binding events would be independent and there would be no activation.

**Figure 6.**
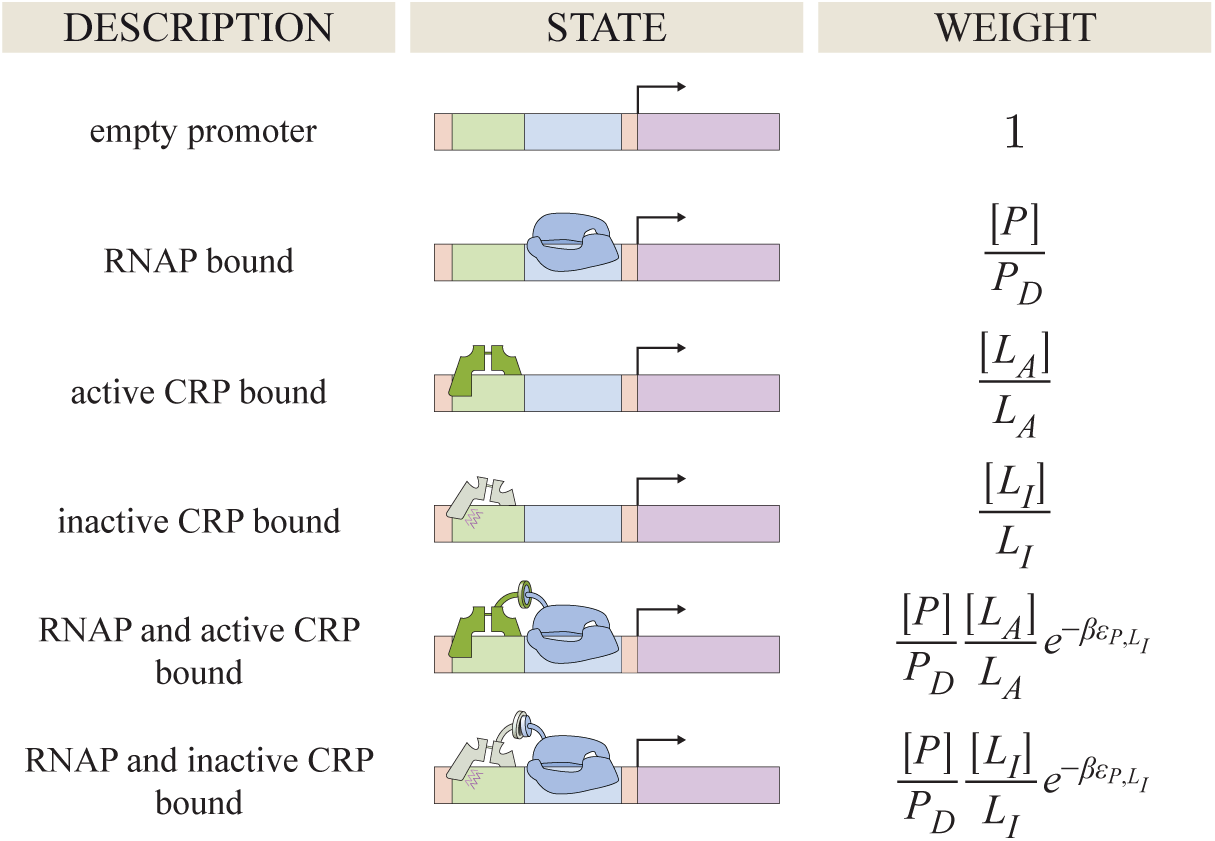
States and weights for a simple activation motif. Binding of RNAP (blue) to a promoter is facilitated by the binding of the activator CRP. Simultaneous binding of RNAP and CRP is facilitated by an interaction energy *є*_*P*,*L_A_*_ for active CRP (dark green) and *є*_*P*,*L_I_*_ for inactive CRP (light green). cAMP (not drawn) influences the concentration of active and inactive CRP as shown in Fig. 4.

We assume that gene expression is equal to the product of the RNAP transcription rate *r*_trans_ and the probability that RNAP is bound to the promoter of interest, namely,

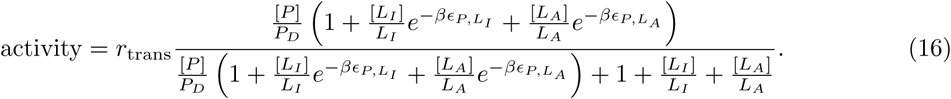

Several additional factors influence gene expression *in vivo*. First, cAMP is synthesized endogenously by *cya* and degraded by *cpdA*, although both of these genes have been knocked out for the data set shown in Fig. 7 (see Methods and Ref. (7)). Furthermore, cAMP is actively transported out of a cell leading to a smaller concentration of intracellular cAMP. Following Kuhlman *et al.*, we will assume that the intracellular cAMP concentration is proportional to the extracellular concentration, namely, *γ*[*M*] (with 0 < *γ* < 1) (31, 32). Hence, the concentration of active CRP satisfies 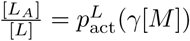 where the fraction of active CRP 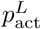 is given by Fig. 2(A) as

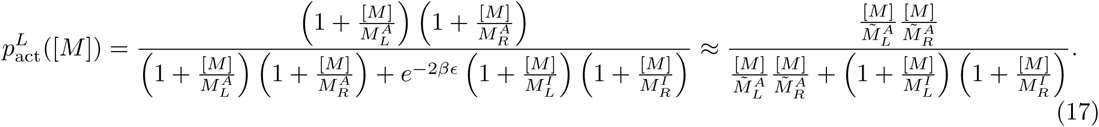

**Figure 7.**
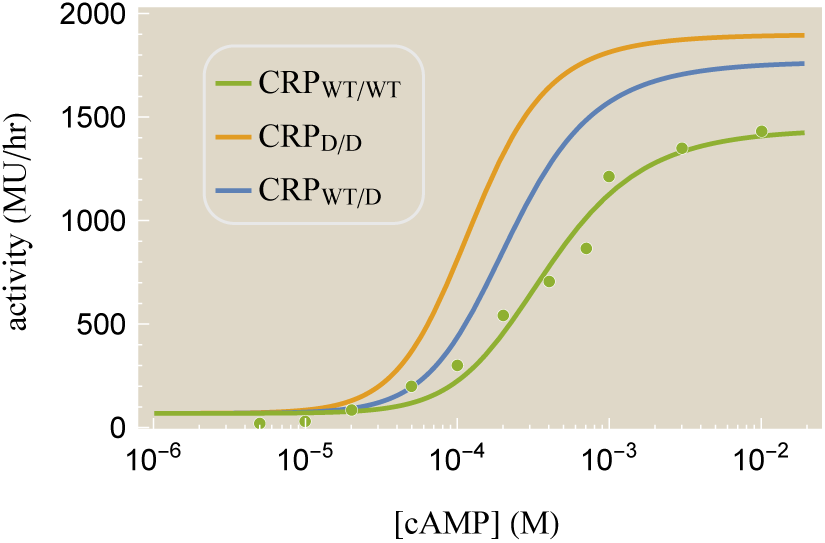
Predicted gene expression profiles for a simple activation architecture. Gene expression for wild type CRP (green dots from Ref. (7)), where 1 Miller Unit (MU) represents a standardized amount of *β*-galactosidase activity. This data was used to determine the relevant parameters in Eq. 16 for the promoter in the presence of [*L*] = 1.5*μ*M of CRP (33). The predicted behavior of the CRP mutants is shown using their corresponding cAMP dissociation constants. Parameters used were 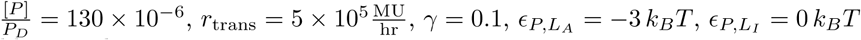, and those shown in Tables 1 and 2.

In the last step, we have again introduced the effective dissociation constants from Eqs. 3 and 4 and dropped any terms proportional to *e*^—*β*_є_^. In addition to these considerations, proteins *in vivo* may experience crowding, additional forms of modification, and competition by other promoters. However, since our primary goal is to understand how CRP mutations will affect gene expression, we proceed with the simplest model and neglect the effects of crowding, modification, and competition.

Because of the uncertainty in the dissociation constant *L_A_* between active CRP and DNA (see Table 2), it is impossible to unambiguously determine the transcription parameters from the single data set for wild type CRP shown in Fig. 7. Instead, we select one possible set of parameters 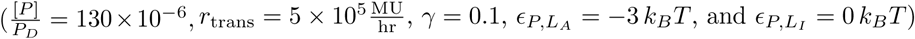 that is consistent with the wild type data. Next, we inserted the other cAMP-CRP dissociation constants (given in Table 1) into Eq. 16 to predict the gene expression profiles of the CRP mutants. Fig. 7 show the possible behavior of the CRP_D/D_ and CRP_WT/D_ mutants. As expected, replacing a WT subunit with a D subunit shifts the gene expression profile leftwards since the D subunit has a higher cAMP affinity (see Fig. 3(A)). Interestingly, the substitution of WT with D subunits comes with a concomitant increase in the maximum gene expression because at saturating cAMP concentrations, a larger fraction of CRP_D/D_ is active compared to CRP_WT/WT_ (96% and 68%, respectively) as seen by using Eq. 17 and the parameters in Table 1. Note that we cannot predict the behavior of any of the CRP mutants with S subunits due to the large uncertainty in 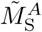.

Lastly, we probe the full spectrum of phenotypes that could arise from the activity function provided in Eq. 16 for any CRP mutant by considering all possible values of the cAMP-CRP dissociation constants 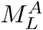, 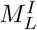, 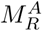, and 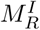 in Eq. 17. In particular, we relax our assumption that cAMP binding promotes the CRP’s active state, as a CRP mutation may exist whose inactive state binds more tightly to cAMP than its active state. Fig. 8 demonstrates that given such a mutation, a variety of novel phenotypes may arise. The standard sigmoidal activation response is achieved when cAMP binding promotes the active state in both CRP subunits 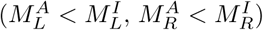. A repression phenotype is achieved in the opposite extreme when cAMP binding favors the inactive CRP state 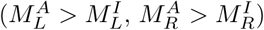. When one subunit is activated and the other is repressed by cAMP 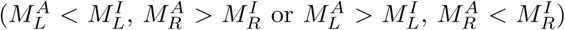, a peaked response can form. If the CRP subunits have the same affinity for cAMP in the active and inactive states 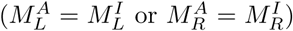, then CRP will behave identically for all concentrations of CRP, generating a flat-line response. It will be interesting to see whether these phenotypes can be achieved experimentally.

**Figure 8.**
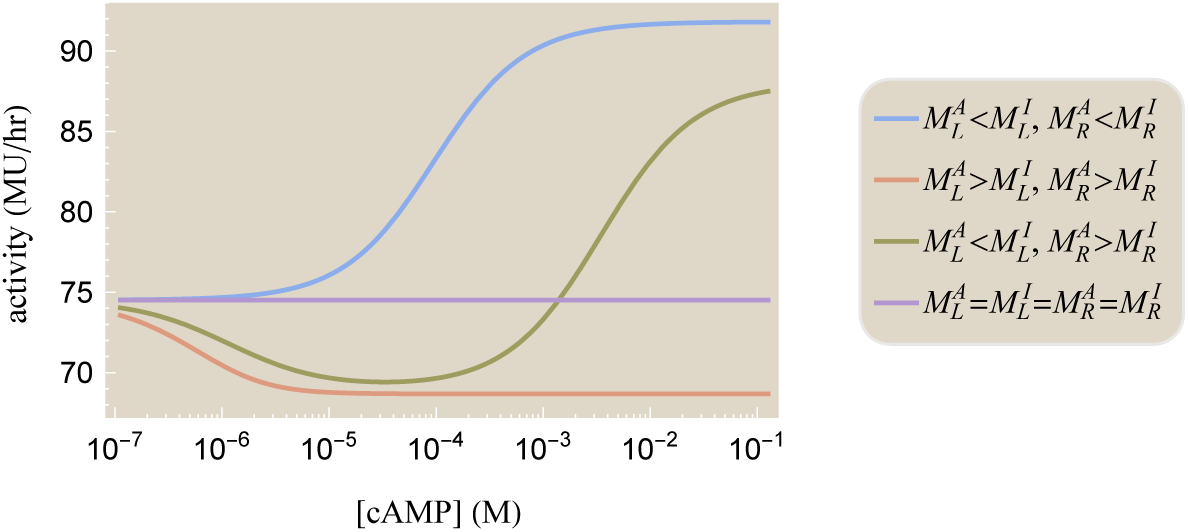
The spectrum of input-output responses for mutant CRP in a simple activation architecture. The possible gene expression profiles given by Eq. 16 can be categorized based upon the cAMP-CRP binding affinity in each subunit. In all cases, we assumed 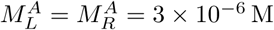. The activation response (blue) was generated using 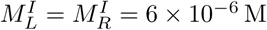. The repression response (orange) used 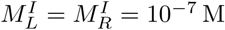. The peaked response (gold) used 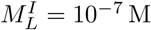 and 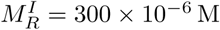. The flat response used 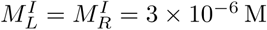. The remaining parameters were the same as in Fig. 7 together with *є* = —3*k_B_T*.

## Conclusion

The recent work of Lanfranco *et al.* provides a window into the different facets of gene regulation through activation (15). Using insights from their *in vitro* experiments, we can break down the process of activation into its key steps, namely: (1) the binding of cAMP to make the activator CRP competent to bind DNA (Fig. 3); (2) the binding of CRP to DNA (Fig. 5); and (3) the recruitment of RNAP to promote gene expression (Fig. 7). By concurrently modeling these processes, we begin to unravel relationships and set strict limits for the binding energies and dissociation constants governing these systems. One hurdle to precisely fixing these values for CRP has been that many different sets of parameters produce the same degenerate responses (see Supporting Information Section A). This parameter degeneracy is surprisingly common when modeling biological systems (34, 35), and we discuss how to account for it within the MWC and KNF models of CRP. A key feature of our analysis is that it permits us to identify the relevant parameter combinations for the system, quantify how well we can infer their values, and suggest which future experiments should be pursued to best constrain the behavior of the system.

Lanfranco *et al.* further explored how mutations in the cAMP binding domain of one or both subunits of CRP would influence its behavior. Specifically, they used three distinct subunits (WT, D, and S) to create the six CRP mutants shown in Fig. 1(B). We analyzed these constructs using both the Monod-Wyman-Changeux and Koshland-Némethy-Filmer models of molecular switching and demonstrated that either model can characterize all six mutants using a self-consistent framework where each subunit is described by a unique cAMP dissociation constant (see Table 1). However, these two models cannot be deemed successful simply because they yield curves that can reproduce the data; the benefit of using mechanistic models involving physical parameters is that the inferred values can be cross-checked against other sources. For instance, the MWC model predicts that singly cAMP-bound CRP will bind more tightly to DNA than unbound or doubly bound CRP, in line with the structural knowledge of the system (10, 18, 28). And while the KNF model can reproduce nearly identical CRP-DNA binding curves to those generated by the MWC model, its inferred parameters imply that CRP should bind equally well to its operator regardless of whether it is singly or doubly bound to cAMP (see Table 2). Overall, these results favor an MWC interpretation of the CRP system.

The models presented here suggest several avenues to further our understanding of CRP. First, several groups have proposed that multiple CRP mutations (K52N, T127, S128, G141K, G141Q, A144T, L148K, H159L from Refs. (9, 36, 37)) only affect the free energy difference *є* between the CRP subunit’s active and inactive states while leaving the cAMP-CRP dissociation constants unchanged. It would be interesting to test the framework developed here across CRP mutants specifically designed to vary these parameters, since the *є* dependence of the system is completely relegated to the effective dissociation constants (see Eqs. 3 and 4).

Second, the MWC and KNF models can be used to predict how the CRP mutants generated by Lanfranco *et al.* would behave *in vivo.* We calibrated the CRP_wt_/_wt_ gene expression profile using data from Ref. (7) and suggested how the remaining CRP mutants may function within a simple activation regulatory architecture given the currently available data (see Fig. 7). It would be interesting to measure such constructs within the cell and test the intersection of our *in vivo* and *in vitro* understanding both in the realm of the multi-step binding events of CRP as well as in quantifying the effects of mutations.

Finally, we note that both the MWC and KNF models can serve as a springboard for more complex descriptions of CRP. It has been debated whether the first cAMP binding event inhibits a second cAMP from binding or if doubly bound CRP has a weaker affinity to DNA (13, 18, 27, 38, 39), and such modifications are straightforward to add to the models discussed above. However, a key advantage of the simple frameworks presented here lies in their ability to *predict* how different CRP subunits combine. For example, in Supplementary Information Section B we demonstrate how the data from the three symmetric CRP mutants in Fig. 3(A) can be used to characterize the asymmetric mutants in Fig. 3(B). It would be interesting to see whether such predictions continue to hold as more mutant subunits are characterized. Such a framework has the potential to harness the combinatorial complexity of oligomeric proteins and presents a possible step towards systematically probing the space of mutations.

## Methods

As described in Ref. (15), the fractional CRP occupancy data in Fig. 3 was measured *in vitro* using 8-anilino-1-naphthalenesulfonic acid (ANS) fluorescence which is triggered by the conformational change of cAMP binding to CRP. The CRP-DNA anisotropy data in Fig. 5 was measured *in vitro* by tagging the end of a 32 bp *lac* promoter with a fluorescein molecule and measuring its anisotropy with a spectrometer. When CRP is bound to DNA, anisotropy arises from two sources: the fast bending of the flanking DNA sequence and the slower rotation of the CRP-DNA complex. Sources of error include oligomerization of CRP, the bending of the flanking DNA, and nonspecific binding of CRP to the DNA.

The *in vivo* gene expression data was taken from Kuhlman et al. using the *lac* operon *E. coli* strain TK310 (7). This strain had two genes knocked out: *cya* (a gene encoding adenylate cyclase, which endogenously synthesizes cAMP) and *cpdA* (encoding cAMP-phosphodiesterase, which degrades cAMP within the cell). Experiments were done at saturating concentrations of inducer ([IPTG] = 1mM) so that Lac repressor negligibly binds to the operator. In this limit, the only transcription factor affecting gene expression is the activator CRP. Gene expression was measured using *β*-galactosidase activity.

## Supporting Material

Supporting Materials with the aforementioned derivations are available online together with a Mathematica notebook that contains all the data, reproduces the fitting (using both nonlinear regression and MCMC), and generates the plots from the paper.

## Author Contributions

TE, JD, and RP performed the research. TE and RP wrote the manuscript.

## Acknowledgements

We thank Lacramioara Bintu for bringing the recent developments on CRP to our attention as well as Terry Hwa, Tom Kuhlman, and Michael Manhart for helpful discussions. All plots were made entirely in *Mathematica* using the CustomTicks package (40) with data obtained from the authors or using WebPlotDigitizer (41). This work was supported in the RP group by La Fondation Pierre-Gilles de Gennes, the Rosen Center at Caltech, and the National Institutes of Health through DP1 0D000217 (Director’s Pioneer Award), R01 GM085286, and 1R35 GM118043-01 (MIRA). We are grateful to the Burroughs-Wellcome Fund for its support of the Physiology Course at the Marine Biological Laboratory, where part of the work on this work was done, and for a post-course research grant (JD).

## Supporting Citations

References (42–44) appear in the Supporting Material.

